# How optimal foragers should respond to habitat changes? On the consequences of habitat conversion

**DOI:** 10.1101/273557

**Authors:** Vincent Calcagno, Frédéric Hamelin, Ludovic Mailleret, Frédéric Grognard

## Abstract

The Marginal Value Theorem (MVT) provides a framework to predict how habitat modifications related to the distribution of resources over patches should impact the realized fitness of individuals and their optimal rate of movement (or patch residence times) across the habitat. Most MVT theory has focused on the consequences of changing the shape of the gain functions in some patches, describing for instance patch enrichment. However an alternative form of habitat modification is habitat conversion, whereby patches are converted from one existing type to another (e.g. closed habitat to open habitat). In such a case the set of gain functions existing in the habitat does not change, only their relative frequencies does. This has received comparatively very little attention in the context of the MVT. Here we analyze mathematically the consequences of habitat conversion under the MVT. We study how realized fitness and the average rate of movement should respond to changes in the frequency distribution of patch-types, and how they should covary. We further compare the response of optimal and non-plastic foragers. We find that the initial pattern of patch-exploitation in a habitat, characterized by the regression slope of patch yields over residence times, can help predict the qualitative responses of fitness and movement rate following habitat conversion. We also find that for some habitat conversion patterns, optimal and non-plastic foragers exhibit qualitatively different responses, and that adaptive foragers can have opposite responses in the early and late phases following habitat conversion. We suggest taking into account behavioral responses may help better understand the ecological consequences of habitat conversion.

## 1 Introduction

In most analyses of the Marginal Value Theorem [5,17], including recent re-analyses [2–4], emphasis is on understanding how changes in the shape of gain functions in patches impacted the optimal residence times and movement rate. Yet an alternative form of habitat alteration is to change the relative frequency of different patch categories, without modifying the categories themselves. For instance one might increase the quality of habitat not by making individual patches richer, but rather by making the richest patches more frequent (and the poorest patches rarer). In the context of the MVT, this corresponds to a situation where the travel time and the gain functions are unchanged, but the frequency distribution of the different patch categories does vary. Whereas changes in the individual gain functions can readily describe scenarios of habitat enrichment [3], modifications of the relative patch frequencies more aptly capture scenarios of habitat conversion (transformation of patches from one category to another). This form of habitat change is illustrated in Fig. 1a. For instance, one might think of converting patches of closed habitat (e.g. forest) into open habitat (e.g. clearings or crop) in a landscape mosaic, or changing the relative frequency of disturbed versus pristine feeding sites [11,16,18]. Alternatively, this can describe the extinction of predators from some patches, or their experimental extirpation, turning hazardous places into risk-free areas [10]. Habitat conversion is actually one pervasive aspect of the current global biodiversity crisis, impacting many different types of ecosystems [8,9,15]. Nonetheless, scenarios of habitat conversion have not received much attention in the MVT literature (see e.g. [14]), probably in part because it requires considering entire habitats, and prevents focusing on individual patches. It further renders classical MVT graphical arguments largely inefficient. Indeed, visualizing optimal residence times as points where the marginal rate of gain equals the long-term average rate of gain 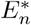; [17]) is of limited help to predict what happens when patch frequencies are modified, as Fig. 1b illustrates.

**Figure 1.**
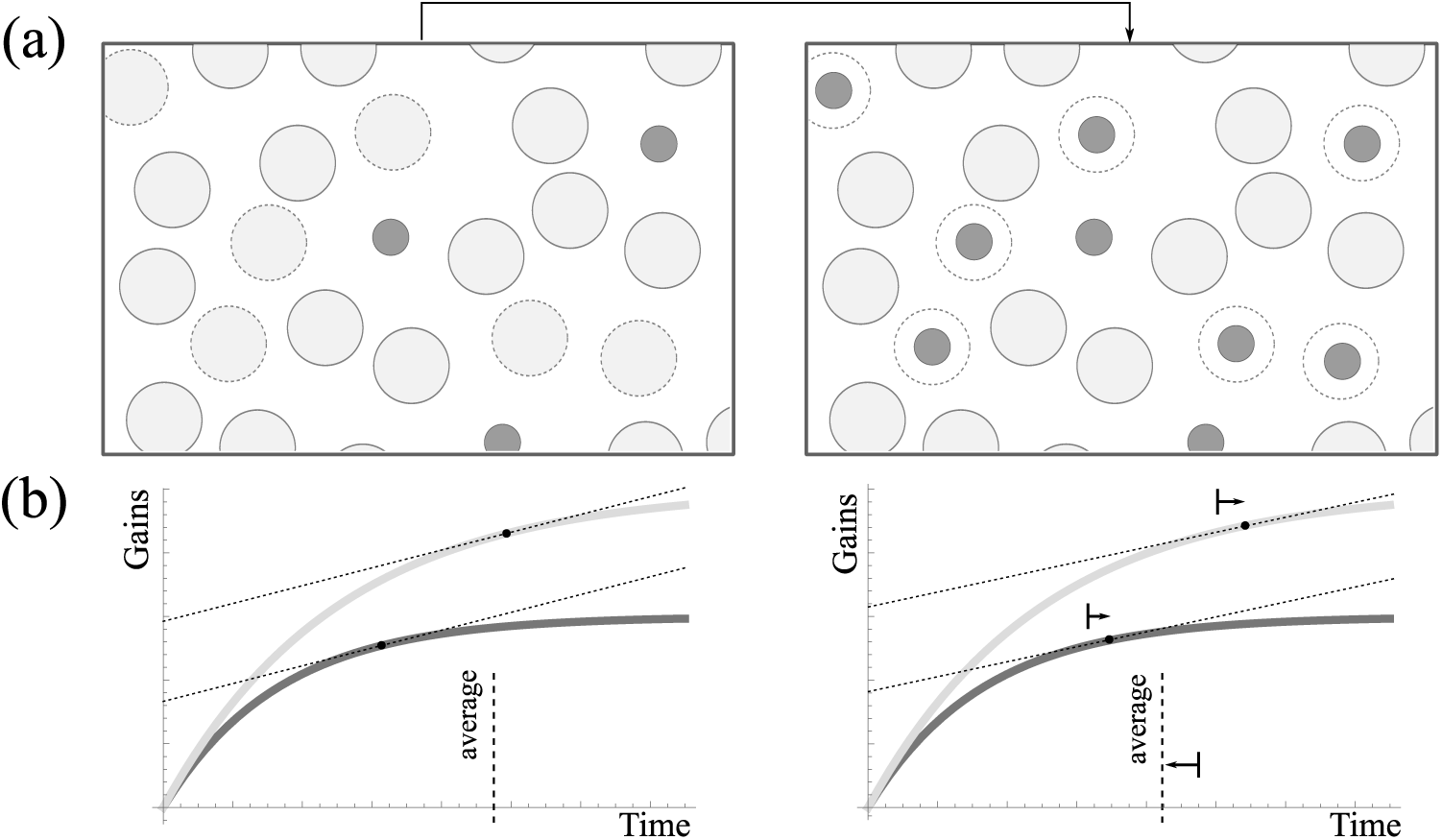
Habitat conversion in the MVT. (a) A patchy habitat with two sorts of patches. Large light disks represent pristine patches, small dark disks disturbed patches. Starting from the situation on the left, with disturbed patches in frequency 1/10, some pristine patches are converted into disturbed patches, resulting in the habitat shown on the right, with 3/10 of disturbed patches. (b) Corresponding changes in residence times under the MVT. It is assumed that pristine patches have more favorable gain functions (light curves) than perturbed patches (dark curves), and the optimal residence times are such that gain functions are tangent to a line of slope
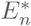 (dotted lines). Increasing the frequency of per-turbed patches decreases 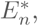 so that the optimal residence time on both patch-types slightly increase. However, the average residence time decreases,and thus the overall movement rate increases, which is difficult to deduce from graphical intuition.

In a previous mathematical reanalysis of the MVT [4], we only briefly mentioned the effect of changing the relative patch frequencies *p_i_* (Section 5 in [4]). We remarked that following changes in the frequency distribution of patch types, all optimal residence times should vary in the same direction, in opposite direction of the long-term realized rate of gain 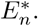 However we did not analyze further how the average residence time 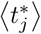and the overall rate of movement in the habitat would respond to such changes in the frequency of different patch types. We simply observed that” *improving habitat quality by manipulating relative patch frequencies decreases all patch residence times*, *i.e. increases the movement rate*”. The last part of the statement is overly simplified. Indeed, the fact that residence time decreases for each individual patch, while true, does not guarantee that the average residence time decreases, since the relative frequencies of different patch types was changed. For instance, if residence times are longer on the best patches, then increasing the frequency of the latter mechanically increases the average residence time, and this might counteract the previous change in behavior observed on any particular patch. This is the kind of scenario illustrated in Fig. 1 (transition from left to right).

One would expect that manipulating the resource distribution over patches through changes in the gain functions [2,3] or through habitat conversion would yield consistent predictions, but this remains to be established. Furthermore, the adaptive response of optimal foragers following habitat change may counteract the direct effects of habitat conversion, and therefore adaptive foragers might in some conditions present qualitatively different responses from static (non-plastic) foragers. To address these points, we here extend our previous reanalyses with a treatment of the effects of changing the relative abundances of different patch-types.

## 2 The MVT and notations

We build on [3,4] and consider the general (heterogeneous) marginal value theorem according to which the optimal residence time 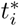 on patches of type *i* is defined implicitly by

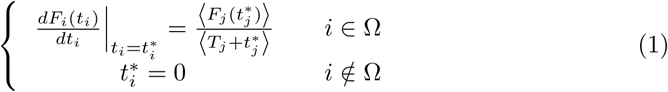

where *F_i_* is the gain function in patches of type *i*, *T_i_* is the average time to reach a patch of type *i* (travel time, usually regarded as the same for all patches) and Ω refers to the set of patches that are effectively exploited in the habitat. Brackets are used to denote spatial averages over the entire habitat (Ω ∪ Ω^**C**^), i.e.

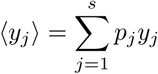

with *p_j_* the frequency of patches of type *j*, for a total of *s* different patch-types (*j* ∈ (1, *s*)).

Quantity
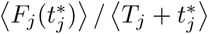
is the long term average rate of gain realized in the habitat (called
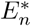 for short). A well-known consequence of eq. (1) is that all exploited patches should be left at the same quitting rate (instantaneous rate of gain at the time the individual leaves) equal to the long term average rate of gain in the habitat 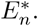

Following [3], we remark that unexploited patches can be gotten rid of by restricting the system to the *s̃* patches that are effectively exploited. This implies rescaling the *p_j_* to new *p͠_j_* accordingly and increasing the travel times to some effective values *T̃_j_*, given by

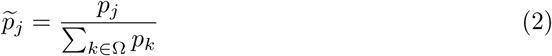

and

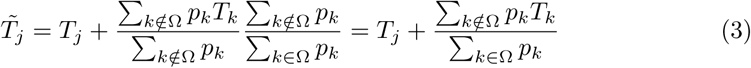

In these conditions, instead of (1), we can study the slightly simpler equation

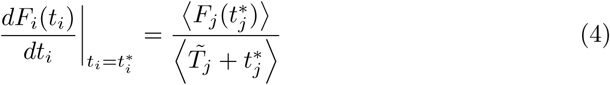

where there are only *s͠* = card(Ω) patch-types to consider, all effectively exploited (i.e. 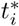 > 0 for all *i*).

For simplicity we will henceforth restrict our attention to the set of patches that are effectively exploited. We will drop the tildes and ignore Ω, but one should remember that *T_j_*, *s* and *p_j_* are intended as their modified values introduced above.

An important quantity to characterize the optimal MVT strategy is the patch-exploitation pattern, quantified by the regression slope (within a given habitat) of patch yields 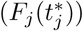 over patch residence times 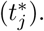 We will call it *ρ_INTRA_* (see [2]). Depending on patch characteristics (shape of the gain functions) and travel time, *ρ_INTRA_* can be positive or negative [2,4], and it will prove important in predicting the consequences of habitat conversion.

## 3 Habitat conversion: manipulating the patch frequency distribution

### 3.1 Which are the best patches?

Differentiating (4) with respect to *p_i_* we get

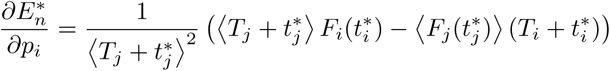

or, isolating 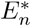:

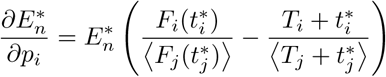

Of course the total variation in 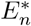 will further depend on how the other patch-types change in frequency; see next Section). Still, requiring the partial derivative to be positive, we get a definition of a “good” (or “better than average”) type of patch as one that, if made more frequent in the habitat, would tend (marginally) to increase the realized fitness 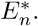 This yields the condition

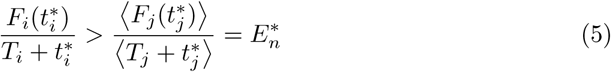

Note that this criterion to be a good patch is close to the criterion to be effectively exploited (i.e. in Ω). Indeed, only patches such that 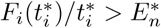 should be exploited and the others should be entirely disregarded [7]. This condition to be an exploited patch is, as expected, less stringent that the condition to be a good patch (remark that the value of *T_i_* does not appear at the denominator, in contrast to eq. (5)).

From (5), the good patches are characterized by a large 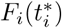 and/or a small 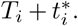 Two scenarios can thus be distinguished regarding the identity of those patches. First, they can have both shorter exploitation time and greater absolute gains, in which case the two effects described above work jointly, and the best patches are unambiguously the best. Alternatively, the best patches can have greater absolute yields, but longer exploitation time, or smaller yield and shorter exploitation time. In this case the two effects work against one another, and the best patches are not the best in all aspects. In this situation their identity will depend quantitatively on the relationship between absolute yield and exploitation time. Examples of the two situations are shown in Fig. 2.

**Figure 2.**
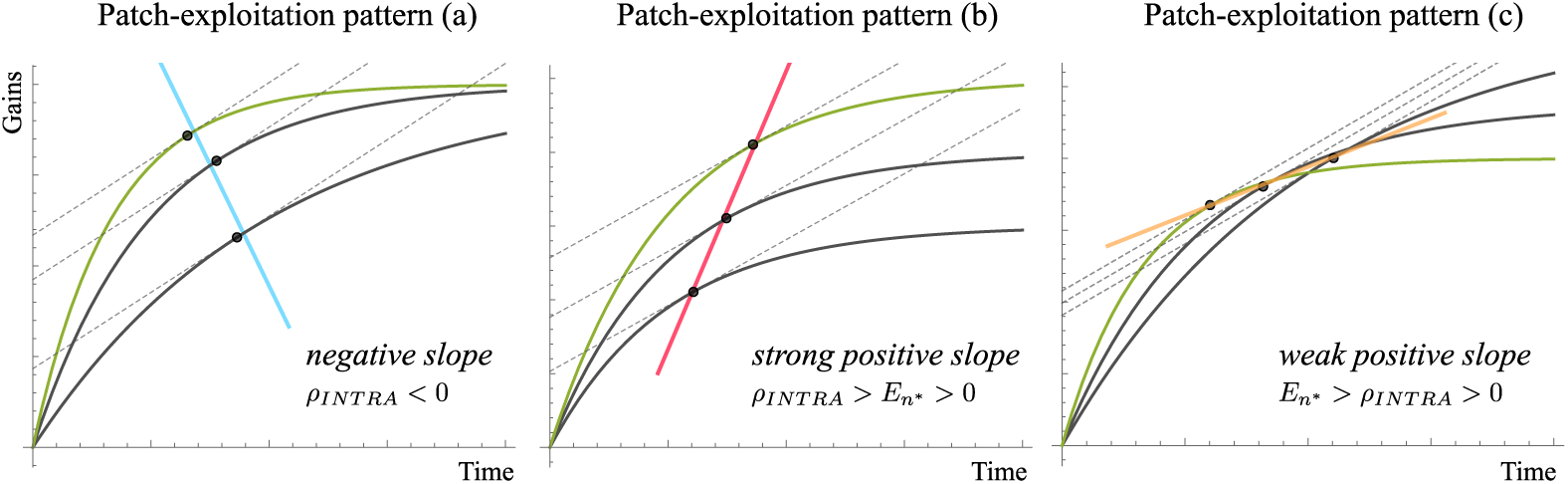
Patch-exploitation patterns within habitats. Hypothetical habitats are shown, with the best patch type (*sensu* eq. 6) in green. Three types of patch-exploitation patterns are useful to distinguish: (a) Best patches are the best in all aspects: they have both greater absolute yield and shorter residence time. This is detectable as a negative association of patch yield and residence time within the habitat (*ρ_INTRA_* < 0). In (b) and (c) the best patches are not the best in all aspects, corresponding to a positive association of patch yield and residence time within the habitat (*ρ_INTRA_* > 0). In (a) they have greater absolute yield but longer residence time, while in (c) they have shorter residence time but lower yield. In (b) the association is strong, i.e. the regression slope of yield over residence time is steeper than the long-term average rate of gain 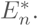 The situation in Fig. 1 also fell into this category. In (c) the association is weak, i.e. *ρ_INTRA_* is shallower than 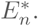 The three situations were obtained with the standard gain function *n*(1 – exp(–*ht*)). In (a) patches differ in resource accessibility (*h*); in (b) they differ in resource content (*n*); and in (c) they differ in both.

### 3.2 Response of realized fitness to habitat conversion

Any particular pattern of habitat conversion will induce changes in at least two of the *P_j_*, while keeping the following constraint satisfied:

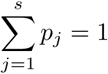

In order to describe the action of habitat conversion, we introduce a variable *x* that encodes the specific pattern of conversion (i.e. which patch-types were converted into which). Changing the value of *x* may alter every patch relative frequency (holding the above constraint satisfied), in a way that describes the type of habitat conversion considered (for example, an increase in the frequency of poor versus good patches). The definition of *x* can be understood as a direction in the space of the *p_j_* along which habitat change operates. We consequently let all *p_j_* be functions of *x*, and the above constraint implies

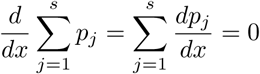

The total variation of 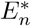 with respect to *x* is thus obtained as the sum of the partial effects of each frequency change, i.e.:

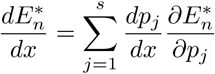

Considering we are at an optimum with respect to all residence times, we can omit any variation in the 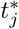 from the derivative of 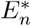 (see [4]), and it follows

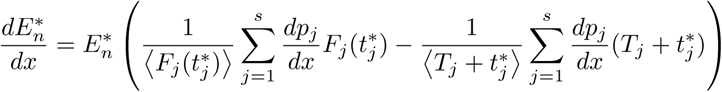

which we may want to rewrite as, in terms of relative variations:

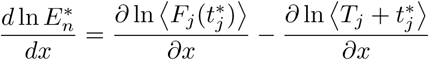

Here, and throughout the rest of the paper, partial derivatives indicate direct variation through the *p_j_*, omitting the indirect variation occurring through changes in the optimal residence times, i.e. *∂/∂x* = Σ *dp_j_*/*dx ∂/∂p_j_*.

Depending on which patch-types are converted into which (i.e. on the habitat conversion pattern *x*), realized fitness may increase or decrease. One might of course want to define ”habitat quality” a priori, irrespective of its actual impact on the realized fitness of an optimal forager. Here we instead take the general (and internally consistent) approach of requiring that habitat quality (at least as perceived by the individual) increases 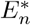 ( [4]). Therefore a specific habitat conversion pattern will be said to increase overall habitat quality if and only if it causes an increase in realized fitness.

From the above equation 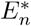 increases with *x* if and only if

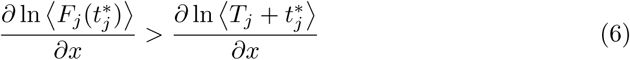

Quite intuitively, this shows that habitat conversion can increase the quality of the habitat in two different ways. First, by making more frequent the patch-types that provide the greatest absolute gains (positive left-hand side in (6)). Second, by making more frequent the patches that do not take long to exploit (those that are easy to reach and/or with short residence times; negative right-hand side). The net effect of habitat conversion on realized fitness rests on the balance of these two effects.

Here, it is important to stress that the variation in realized fitness of an optimal forager is not influenced by the plastic response of residence times (see also [4]). It follows that our definitions of patch and habitat quality, and thus the response of fitness to habitat conversion, are equally valid for optimal (perfectly plastic) or static (non-plastic) foragers, i.e. those that would keep their patch-exploitation pattern unchanged following habitat conversion.

Comparing eq. (6) with the condition for 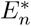 to increase when manipulating patch characteristics (equation (6) in [4]), we see that it is almost identical: the first term is the same, and the second is similar, only lacking the penalty term 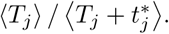 This is because when varying *T_j_* as a patch attribute, we only operate on a fraction of the entire time it takes to exploit one patch (while the optimal residence time itself was out of external control). Here, in contrast, when converting patches from one type to another, the entire duration of the exploitation cycle (travel and residence time) is impacted, and the penalty factor therefore vanishes to unity.

Remark that since our sensitivity analysis approach considers infinitesimal changes in the *p_j_*, conclusions are unaffected by the implicit treatment of unexploited patches (see (2) and (3) above). Indeed, infinitesimal changes generically do not change Ω, and the latter can be regarded as a constant. Of course, to predict the consequences of sustained changes (i.e. to integrate over *x*), one should remember to update Ω appropriately, when a patch-type leaves or enters the set of exploited patches.

### 3.3 Response of average movement rate to habitat conversion

We now consider the effect of habitat conversion on the optimal average movement rate. In the context of the MVT, the latter is defined as the rate of patch switching 1/ 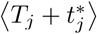, i.e. the inverse of the average time needed to exploit one patch, following [3]. For a systematic (directional) forager, this rate of patch-switching controls the linear speed through the habitat [1], and for a random-walker it sets the timescale of the exploration and is as such expected to be proportional to the diffusion coefficient [20]. In what follows we will thus study the variation of 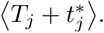 This quantity is closely related to another quantity of interest in the field of behavioral ecology, the average residence time 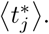

Obviously, the total variation in 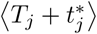 can be split into two components. First, there is the direct effect of changing the relative frequency of the different patches, which mechanically alters the average. This effect is

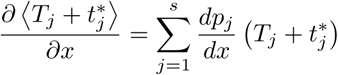

Second, there is an indirect effect whereby the resulting change in 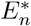 (computed in the previous section) impacts all individual optimal residence times 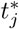 (but not travel times, obviously). It follows that

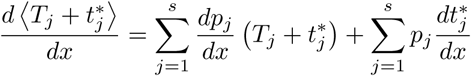

Replacing the total variations of optimal residence times with their expression (see [4]), some calculations (Appendix) yield the following criterion for the average movement rate to increase with *x*:

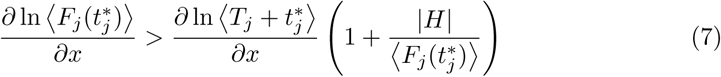

where, as in [4], *H* is the harmonic mean of the second time derivatives of gain functions (*H* < 0).

This is very similar to eq (6), except that the relative change in patch exploitation time (right-hand side) is multiplied by a constant greater than one (parenthesis). The latter depends on the shape of the gain functions (height and second time-derivative). Therefore the impact of the change in patch exploitation time is amplified compared to eq. (6).

The previous result holds true for an optimal forager that has quickly enough adjusted its movement strategy following habitat conversion. If, on the extreme opposite, we consider a non-plastic forager (or, similarly, a plastic forager before it had time to update its strategy), the observed indirect variation of optimal residence times 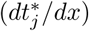would be nil, and the criterion for average movement rate to increase is then simply:

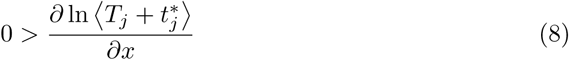

## 4 Discussion

### 4.1 Consequences of habitat conversion for fitness and movement rate

Combining (6) and (7), we obtain a simple graphical representation of what impact any pattern of habitat conversion would have on fitness and movement rate for an optimal forager, and from (6) and (8) the corresponding for a non-plastic forager (Figure 3). All conversion patterns can be sufficiently characterized with a pair of quantities: the relative change in average patch-exploitation time 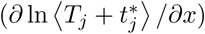 and the relative change in patch yield 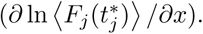 This reveals some constraints on the possible patterns of co-variation of fitness and movement rate following habitat conversion.

**Figure 3.**
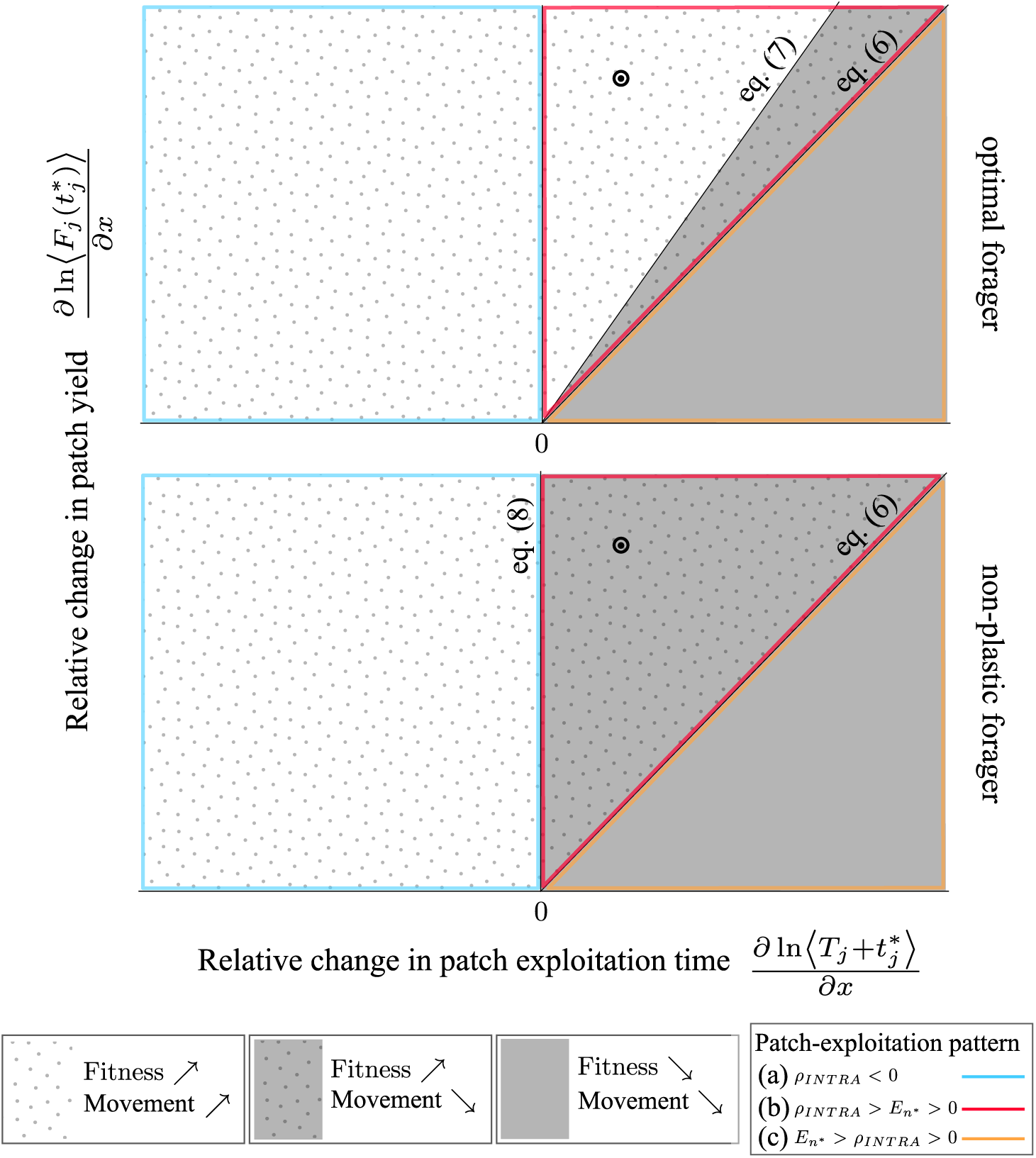
Predicting the impacts of habitat conversion: a summary. The graph summarizes mathematical predictions on the effect of habitat conversion on realized fitness and on average movement rate, for an optimal forager (*top*) or a non-plastic forager (*bottom*). The plane shown represents all possible scenarios of habitat conversion, as characterized by *∂*ln 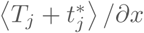 (x-axis) and *∂* ln 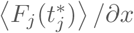 (y-axis). Three areas can be identified that yield qualitatively different predictions (legend on the left). Note that fitness and movement rate can covary negatively only in the gray dotted area, that is much smaller for an optimal forager. The pattern of patch-exploitation prior to habitat conversion (see Figure 2) can be used to determine in which part of the plane we are in (see legend). Circled points locate a specific scenario of habitat conversion that will be simulated numerically in Fig. 4.

**Figure 4.**
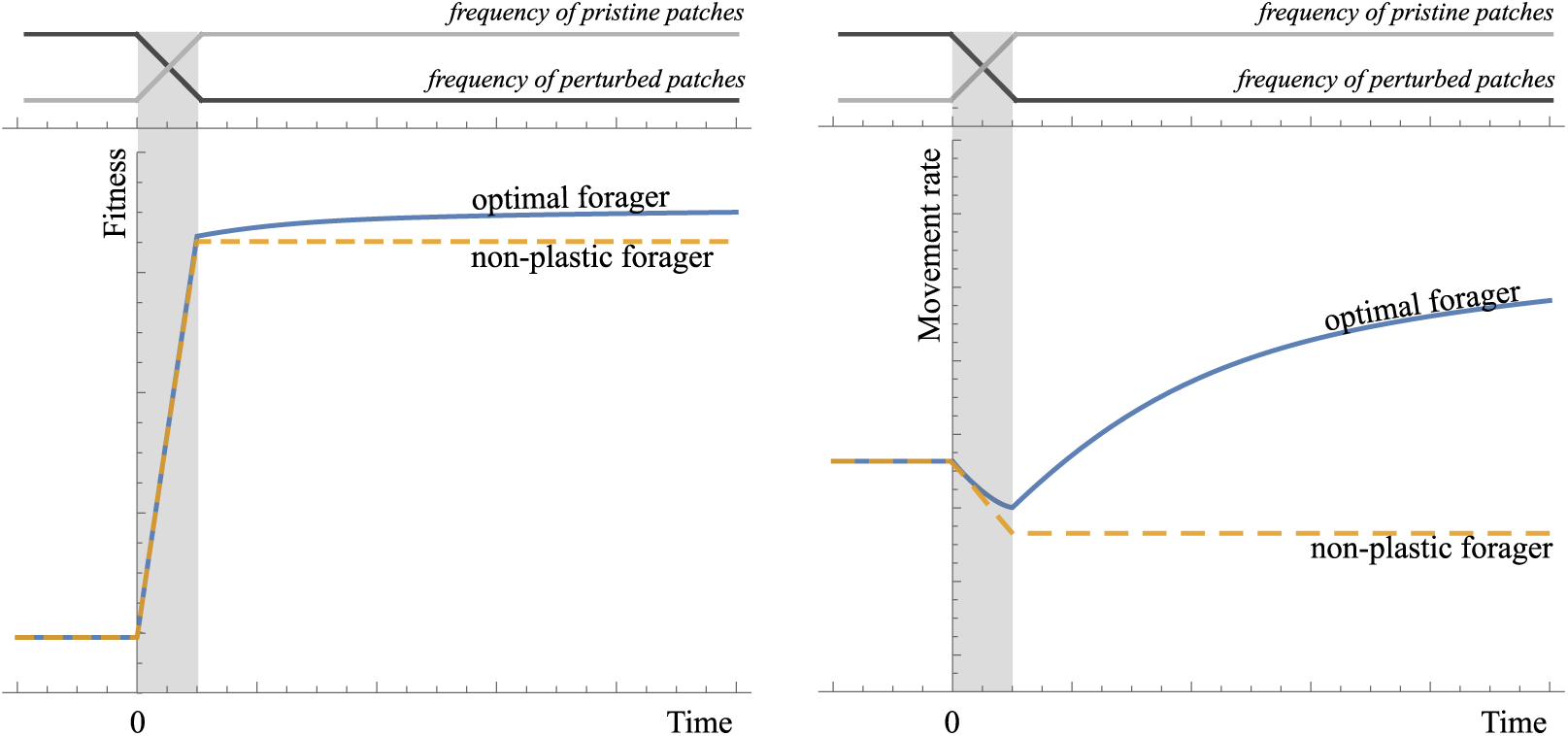
A simulated example of habitat conversion. In a two patch-type habitat (pristine versus perturbed patches; same gain functions as in Fig. 1 & 2), between time 0 and time 1 (shaded parts) the frequency of pristine patches is gradually increased from 1/5 to 3/4. The variation of fitness (left) and average movement rate (right) are shown through time for two foragers: an optimal forager that gradually adjusts its patch-exploitation pattern to the new habitat conditions, and a non-plastic forager, that maintains the same patch exploitation pattern. Note that the two foragers initially present the same qualitative responses (negative covariation of fitness and movement), but ultimately qualitatively different responses (the optimal forager adopting greater movement rate). In our simulation habitat conversion caused a 61% increase in 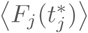 and a 3% increase in 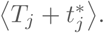 As an indication these values were located as a circle on Fig. 3; simulation results conform to mathematical predictions.

First, we can see that for all types of foragers, plastic or non-plastic, habitat conversion patterns that decrease habitat quality (realized fitness) will necessarily decrease average movement rate as well (gray area). Conversely, if habitat conversion results in higher average movement rate, then it should increase overall fitness as well (white dotted area). In other words, if fitness decreases or average movement rate increases, fitness and movement necessarily co-vary positively. However, there is the possibility of a negative covariation for some habitat conversion patterns (gray dotted area).

Importantly, the likelihood of a negative co-variation of fitness and movement is much more important for a non-plastic forager compared to an optimal forager (the white dotted area in larger in Fig. 3). For optimal foragers, a negative covariation of fitness and movement rate only occurs for a narrow set of habitat conversion patterns (gray dotted area in Fig. 3), feasible only if

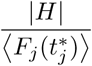

is large enough. In graphical terms, this means that the gain functions should be sufficiently curved (concave), and consistently so (*H* is a harmonic mean, very sensitive to low values), relative to their height.

As a consequence, there is an entire set of habitat conversion patterns, comprised between the lines of eq. (7) and eq. (8) in Figure 3, for which optimal foragers and non-plastic foragers will exhibit qualitatively different responses. This difference between optimal and non-plastic foragers would also manifest itself as a difference between the initial and final responses of an optimal forager. Indeed, following habitat conversion, an adaptive foraging that gradually (rather than instantaneously) adjusts its patch-exploitation pattern [12], would initially exhibit a negative covariation of fitness and movement rate, but ultimately reverse to a positive covariation, once it has fully updated its strategy. Numerical examples of these situations (see Appendix for simulation methods) are provided in Figure 4.

### 4.2 Predicting the consequences of habitat conversion from the initial pattern of patch exploitation

We can obtain further predictions under the reasonable and common assumption that all patches have on average the same travel time *T*, or equivalently that travel times show no consistent response to habitat conversion (see Appendix). This occurs for instance if travel times are controlled by the spatial location of patches. In these conditions optimal movement rate is entirely governed by average optimal residence time, and the two are just inversely related [3].

In this case, the partial derivative on the right-hand side of eq. (6), (7) and (8) is entirely controlled by the residence times on converted patches. It is therefore possible to discriminate different portions of Figure 3 (i.e. different possible habitat conversion patterns) from the observed pattern of patch exploitation in the (unperturbed) habitat (Fig. 2). In practice, one must compute the regression slope of patch absolute yields 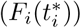 over residence times 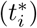 in the initial habitat, that we called *ρ_INTRA_* ( [2]). This regression slope provides the generic value of 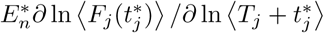 following habitat conversion (an exact value with two patch-types, and a value representative of most possible habitat conversion patterns in the general case). It follows that the three contrasted patch-exploitation patterns shown in Figure 2, discriminable from the regression slope *ρ_INTRA_*, generically produce habitat conversion patterns that map onto different parts of Figure 3.

On Fig. 3, one can see that patch-exploitation pattern (a), i.e. a negative regression of patch yield over residence time, ensures a positive covariation of fitness and movement following habitat conversion, for optimal or non-plastic foragers alike. The same is true for patch-exploitation pattern (c), except that changes will have the opposite sign. However, for patch-exploitation pattern (b) (strong positive regression slope), whilst a negative co-variation is ensured for non-plastic foragers, no firm prediction can be achieved for an optimal forager(Fig. 3). Indeed, prediction further requires knowledge of which patches were converted, and of the curvature of fitness functions (parameter *H* in eq. 7), to tell apart the gray and white dotted areas. Furthermore, it is for patch-exploitation pattern (b) that one would expect qualitative differences between optimal and non-plastic foragers, or non-monotonous dynamical responses from adaptive foragers (see Figs. 3 & 4).

### 4.3 Conclusions

We envisioned habitat change in the Marginal Value Theorem from the perspective of habitat conversion, i.e. changes in the relative frequencies of the different categories of patches. This differs from the classical approach of varying the shape of the gain functions [3,4]. The predictions obtained are in good agreement with previous re-analyses of the MVT [2,4]. For example:

- An increase in habitat quality may have any (negative, null or positive) impact on average movement rate (or average residence time).
- An increase in average movement rate with habitat quality requires less stringent conditions [2,4]. While a negative correlation of patch yield and residence time ensures that an increase in habitat quality will increase average movement (patch-exploitation pattern (a); leftmost halves of Fig. 3), a positive correlation does not guarantee a decrease in movement rate.
- For an optimal forager, if there is a strong positive correlation of patch yield and residence time (pattern (b)), one cannot predict the response of average movement to habitat quality based solely on simple average quantities (average patch yield, average residence time, average rate of gain 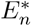). We further need to know the harmonic mean of the gain function second-time derivatives (|*H*|; see eq. (7)). The greater |*H*|, i.e. the more curved the gain functions, the more likely it is to observe a decrease in movement with habitat quality. Consistent results were obtained with the other form of habitat change (see Theorem 3 and eq. (17) in [4]).

It thus seems one can safely transpose results obtained with one approach to the other. We can expect existing predictions regarding the consequences of changing the resource distribution in a habitat that were derived from changing the gain functions [2–4,6,21] to extend to scenarios of habitat conversion as well. This also suggests one might obtain an integrated mathematical treatment of the consequences of habitat changes under the MVT.

In this work we also obtained novel predictions that can guide our prediction and interpretation of actual responses to habitat conversion. For instance, the pattern of patch-exploitation in the pristine habitat, that can be characterized from a simple regression line of of patch yields over patch residence times (*ρ_INTRA_*; Fig. 2; see also [2]), can be used to predict what type of responses to expect following habitat conversion (Fig. 3). Furthermore, only some types of habitat conversion patterns, occurring for patch-exploitation pattern (b), are expected to generate qualitatively different responses for plastic and non-plastic foragers. Last, adaptive foragers could for some habitat conversion patterns exhibit qualitatively different responses in the short and in the long-term (Fig. 4), making it challenging to extrapolate ultimate consequences from initial responses.

While the theoretical consequences of habitat conversion are usually considered at the scale of populations and communities [13,19], we suggest the MVT can offer a framework to apprehend these questions at the individual and behavioral scales. For instance, dynamic changes in the movement strategies of individuals on the short-term may change the relative exploitation of different parts of the habitat, as well as the realized level of migration and connectivity. Depending on their sign, behavioral responses might mitigate, or on the contrary amplify, the population and community responses expected under the assumption of static migration rates and connectivity levels. A better integration of the behavioral and population scales might therefore improve our ability to predict the ecological consequences of habitat change in general, and habitat conversion in particular.

## Footnotes

**Conflict of interest disclosure** The authors of this preprint declare that they have no financial conflict of interest with the content of this article.

## Acknowledgments

This work was supported by INRA and Université Côte d’Azur (IDEX JEDI). This preprint has been reviewed and recommended by Peer Community In Ecology (https://dx.doi.org/10.24072/pci.ecology.10005)

## Appendix Derivation of eq. (7)

As explained in Section 3.3 in the main text, and following [4] to express the indirect variation of 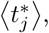 we can formulate the total variation of 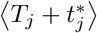 as :

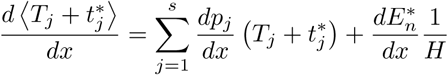

where *H* is the harmonic mean of the second time-derivatives, i.e.

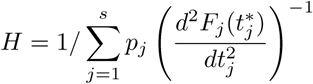

Replacing the variation of 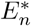 with its expression (given in Section 3.2 in the main text), we get

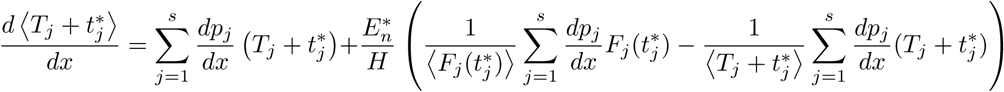

We can directly identify the partial log derivative of 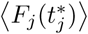 and 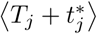 in the parenthesis, and upon factoring 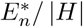 out we obtain:

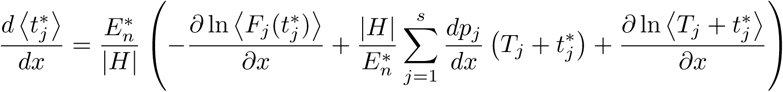

Introducing the partial log derivative of 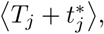 and requiring the parenthesis to be negative (since average movement rate varies in opposite direction of 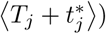 yields

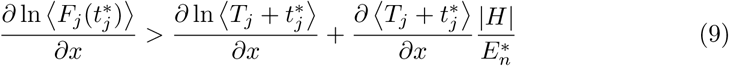

This, replacing 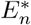 with its expression, yields eq. (7).

Now, if all patches have on average the same travel time *T*, or if variation in travel time shows no consistent trend with habitat conversion, we have *d* 〈*T_j_*〉/*dx* = 0. Note that under the second scenario (i.e. if all travel times are not equal), achieving a null derivative would in practice requires specific forms of habitat conversion, owing to the constraint that the sum of the *p_j_* is constant. The constraint gradually vanishes as the number of patch-types (*s*) gets large. From *d* 〈*T_j_*〉/*dx* = 0 it follows:

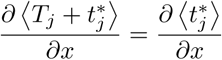

### Numerical simulations

In order to generate the numerical simulations presented in Figure (4), we used a simple gradient ascent algorithm. Individuals were assumed to update each residence time (*t_j_*, omitting the asterisk as they need not be at optimal value) gradually, in the direction that (locally) increases the long-term average rate of gain 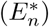 and at a rate proportional to the fitness differential, i.e.

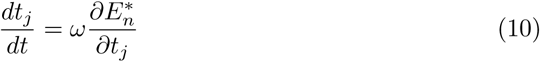

where *t* denotes ecological time (on which habitat changes take place) and *ω* is a constant quantifying the speed of behavioral adjustments.

Habitat conversion is modeled by specifying how the patch relative frequencies change through time, i.e. by specifying functions *p_j_*(*t*). In the simulations of Fig. 4, the function were taken to be linear, so that relative frequencies changed linearly from their initial to their final values. It was assumed that individuals, prior to the onset of habitat change, had settled at the optimal residence times for the initial habitat. When *ω* is very large the forager is effectively optimal and immediately adjusts its strategy to match current habitat conditions (in accordance with the MVT). When *ω* = 0 the individual does not adjust its strategy (non-plastic forager). Intermediate values of *ω* represent less-than-perfect plastic foragers that gradually adapt to habitat changes.

